# ProteomeVis: a web app for exploration of protein properties from structure to sequence evolution across organisms’ proteomes

**DOI:** 10.1101/227827

**Authors:** Amy I. Gilson, Niamh Durfee, Rostam M. Razban, Hendrick Strobelt, Kasper Dinkla, Jeong-Mo Choi, Hanspeter Pfister, Eugene Shakhnovich

**Author notes:** These authors contributed equally (Amy I. Gilson, Niamh Durfee, Rostam M. Razban).

## Abstract

**Motivation:** Protein evolution spans time scales and its effects span the length of an organism. A web app named ProteomeVis is developed to provide a comprehensive view of protein evolution in the S. *cerevisiae* and *E. coli* proteomes. ProteomeVis interactively creates protein chain graphs, where edges between nodes represent structure and sequence similarities within user-defined ranges, to study the long time scale effects of protein structure evolution. The short time scale effects of protein sequence evolution is studied by sequence evolutionary rate (ER) correlation analyses with protein properties that span from the molecular to the organismal level.

**Results:** We demonstrate the utility and versatility of ProteomeVis by investigating the distribution of edges per node in organismal protein chain universe graphs (oPCUGs) and putative ER determinants. S. *cerevisiae* and E. *coli* oPCUGs are scale-free with scaling constants of 1.78 and 1.50, respectively. Both scaling constants can be explained by a previously reported theoretical model describing protein structure evolution (Dokholyan *et al.,* 2002). Protein abundance most strongly correlates with ER among properties in ProteomeVis, with Spearman correlations of −0.51 (p-value<10^−10^) and −0.46 (p-value<10^−10^) for S. *cerevisiae* and *E. coli*, respectively. This result is consistent with previous reports that found protein expression to be the most important ER determinant (Zhang and Yang, 2015).

**Availability:** ProteomeVis is freely accessible at http://proteomevis.chem.harvard.edu/proteomevis/.

**Supplementary information:** Available to download on bioRxiv.

## 1 Introduction

Protein evolution spans time scales, from the billions of years during which the universe of extant protein structures emerged to the fraction of microseconds in which protein sequences mutate (Caetano-Anollés *et al.*, 2009). Protein evolution also spans length scales, from a protein’s length and stability, to its copy number and interactions with other proteins, and to its influence on organismal fitness (Zeldovich and Shakhnovich, 2008; Sikosek and Chan, 2014; Bershtein *et al.*, 2017). Accordingly, protein evolution is studied through lines of inquiry usually separated by scale.

One approach takes a long time scale view, inferring evolutionary dynamics of proteins by ordering extant proteins into a graph based on relationships between their structures and sequences (Qian *et al.*, 2001; Dokholyan *et al.*, 2002; Koonin et al., 2002; Deeds *et al.*, 2004). Dokholyan *et al.*, 2002 constructed the graph by representing protein domains as nodes and connecting pairs of nodes with edges only if their structure similarities, measured by DALI Z-scores (Holm and Sander, 1993), exceed 9 and their sequence identities (SIDs) are less than 0.25. A Z-score of 9 was found to differentiate protein pairs with similar (Z-score>9) or different (Z-score<9) structures (Dokholyan *et al.*, 2002). A SID of 0.25 originates from numerous studies finding that two aligned protein sequences sharing greater than a SID value ranging from 0.2 to 0.4, are likely to share the same structure (Chothia and Lesk, 1986; Sander and Schneider, 1991; Chung and Subbiah, 1996; Rost, 1999). Such sequence pairs are called homologous.

Strikingly, the distribution of edges per node in this non-homologous protein domain graph, dubbed the protein domain universe graph (PDUG), scales as *k*^−1^·^6^, where *k* stands for degree (number of edges per node) and 1.6 is the scaling constant (Dokholyan *et al.*, 2002). Scale-free graphs were first shown to form by a process of preferential attachment, but models of this type predict scaling constants greater than 2 (Albert and Barabási, 2000; Krapivsky *et al.*, 2000; Albert and Barabási, 2002). For the PDUG, an alternative model called Big Bang was developed, where structures originate from gene duplication events followed by significant divergence in sequence. Simulations of the Big Bang model generated scale-free degree distributions with scaling constants ranging from 1.4-1.9. The overall average over 100 independent simulations yielded a scaling constant of 1.6, matching that observed in nature (Dokholyan *et al.*, 2002). Further studies found PDUG sub-graphs built from proteins belonging to 59 prokaryotic organisms’ proteomes are also scale-free (Deeds *et al*., 2004; Roland and Shakhnovich, 2007). These sub-graphs, called organismal PDUGs (oPDUGs), have scaling constants ranging from 1.3- 1.9, closely matching the extent of the Big Bang model (Deeds *et al.*, 2004). The Big Bang model’s successes in predicting scaling constants, indicate that the universe of extant protein structures emerged by a process of divergent evolution (Deeds and Shakhnovich, 2010).

A complementary approach to protein evolution takes a shorter time scale view, inferring determinants of sequence evolutionary rates (ER) by correlating ER with other protein properties (Zhang and Yang, 2015). A complex cellular environment determines the fitness effects of mutations that arise randomly in genes. Diverse and sometimes countervailing selection pressures channel the course of a protein’s evolution (Zeldovich and Shakhnovich, 2008; Sikosek and Chan, 2014; Bershtein *et al.*, 2017). Over the years in which organisms diverge from their common ancestor, selection pressures aggregate in each protein’s ER. ERs span three orders of magnitude (Zhang and Yang, 2015) and by examining which properties correlate with ER, previous work attempted to extract the major constraints of protein evolution.

Protein expression was found to be the most prominent constraint. Quantified by mRNA abundance because it was more tractable to measure than protein abundance (Vogel and Marcotte, 2012), protein expression correlates negatively with ER across all three kingdoms of life, with correlation coefficients exceeding those observed for other quantities (Pál *et al.*, 2001; Drummond *et al.*, 2005, 2006; Drummond and Wilke, 2008; Zhang and Yang, 2015). Much effort has been expended to identify other protein properties that correlate with ER, independent of protein expression. Protein length and contact density were found to correlate positively with ER in *E. coli, S. cerevisiae* and *D. melanogaster* (Bloom *et al.*, 2006; Zhou *et al.*, 2008), while the number of protein-protein interaction partners (PPI degree) was reported to correlate negatively with ER in *S. cerevisiae* (Fraser *et al.*, 2003). However, the evidence is not incontrovertible. In *S. cerevisiae*, a significant negative correlation was found between contact density and ER (Shakhnovich, 2006), and no correlation was found between PPI degree and ER when carefully controlling for protein expression (Bloom and Adami, 2003).

By juxtaposing protein structure graphs and ER correlation analyses, insights into one time scale of protein evolution can be gained by information from the other. For example, not only do proteins with high contact densities tend to evolve more quickly at the sequence level than their less compact counterparts according to Zhou *et al.*, 2008, they also tend to have larger structure neighborhoods (Shakhnovich *et al*., 2005), defined as the number of different sequences encoding a structure. This is because selection for protein stability, roughly approximated by contact density, biases evolution towards exploring closely related structures (Gilson *et al.*, 2017). An example that demonstrates long time scale structural effects explaining sequence evolution constraints: proteins with related structures (Lukatsky *et al.*, 2007) or sequences (Wright *et al.*, 2005) have a statistically enhanced likelihood of forming stable PPIs. This is most evident by the prevalence of homodimers and superfamily of dimers in nature (Ispolatov *et al.*, 2005).

In this work, protein chain graphs are merged with correlation analyses of ER determinants into an interactive visualization tool called ProteomeVis. ProteomeVis’ graph-theoretical approach takes a global view of protein evolution over long time scales to capture structure evolution, while ER correlation analyses span relatively short time scales. ProteomeVis joins the growing list of visualization tools developed to facilitate understanding of complex and interwoven experimental data (Vizcaíno *et al.*, 2015). What makes ProteomeVis unique is the disparate data it integrates. A tool already exists that enables users to interactively create protein graphs based on sequence and structure similarities (Nepomnyachiy *et al.*, 2015). However, no tool, to the best of our knowledge, includes protein properties from the molecular to the organismal level, as well as sequence and structure similarities to create protein graphs to provide a comprehensive overview of protein evolution.

The roadmap for the remainder of the text is as follows. After describing how we collect the underlying data and incorporate it in the web app (Methods), we demonstrate the utility of ProteomeVis by recapitulating and expanding upon previous literature results regarding protein universe graphs and ER determinants (Results). The Results section highlights ProteomeVis’ versatility, illustrating how it facilitates evolutionary studies of protein structure and sequence.

## 2 Methods

### 2.1 Data curation

To access the in-house Python scripts that manage ProteomeVis’ data and create the web app’s static files, please refer to https://github.com/rrazban/proteomevis_scripts. ProteomeVis currently contains *Saccharomyces cerevisiae (S. cerevisiae)* and *Escherichia coli (E. coli*) proteomes. These two organisms have the largest number of protein structures deposited in the Protein Data Bank (PDB) (Berman *et al*., 2000), accessed in November 2017, per proteome size. In addition, *S. cerevisiae* and *E. coli* have a plethora of experimentally measured protein property data since they are model organisms (Cooper, 2000). Future iterations of ProteomeVis will include more proteomes, such as those from *Homo sapiens* and *Mus musculus.*

Data are collected at the level of protein chains. Although most structure evolution studies and some sequence evolution studies analyze protein domains, the advantage of chains is that they correspond directly to genes. Protein chain identification is straightforward, while protein domain identification is dependent on the subjective definition chosen (Ingolfsson and Yona, 2008). In addition, the protein chain is the smallest level at which cellular-wide protein properties such as abundance, protein-protein interaction and dosage tolerance can be measured. For the remainder of the text, the word “protein” is used to mean “protein chain”.

#### 2.1.1 Protein structure identification

The *S. cerevisiae* S288c proteome has 6049 proteins and the *E. coli* K-12 proteome has 4306, manually verified by the Universal Protein Resource (UniProt) (The UniProt Consortium, 2016) and last modified on February 19, 2017 and October 1, 2017, respectively. We develop a procedure that draws upon existing resources to match *S. cerevisiae* S288c and *E. coli* K-12 sequences in UniProt with *S. cerevisiae* and *E. coli* structures in the PDB. First, the Structure Integration with Function, Taxonomy and Sequences resource (SIFTS) (Velankar *et al*., 2013) is employed to identify protein sequences with potentially matching structures. Briefly, SIFTS assigns PDB structures to UniProt sequences with a minimum SID of 0.9 and the source organism being zero, one or two levels up the species level in the taxonomy tree (Velankar *et al.*, 2013). If multiple structures are found for a given sequence, the structure with the largest sequence coverage is chosen. If multiple structures have the largest coverage, the structure solved at the lowest resolution is chosen. Second, protein structures are downloaded from the PDB and their lengths are calculated. Structures are kept if their lengths are at least 80% of that listed for protein chains in UniProt. This step filters out those protein structures that align well with only a small portion of the protein chain, e.g. a single domain from a multidomain protein.

Running through the outlined steps yields 747 *S. cerevisiae* and 1,098 *E. coli* proteins. ProteomeVis includes protein property data for only these proteins. Monthly updates are performed to incorporate modifications to reported proteomes in UniProt and additional structures deposited in the PDB.

#### 2.1.2 Structure similarity and sequence identity

The template modeling score (TM-score) is used as a metric for structure similarity. TM-score and SID for all intra-organism protein pairs of *S. cerevisiae* and *E. coli* are calculated by the TM-align software (Zhang and Skolnick, 2005), freely downloadable from Yang Zhang’s Research Group website. Briefly, TM-align iteratively aligns the two protein structures until its TM-score is maximized and remains stable between iterations (Zhang and Skolnick, 2005). Both TM-score and SID vary from 0 to 1, where 0 signifies complete inequality and 1 signifies identity. TM-score is proportional to the sum over aligned residue pair distance-dependent weights. TM-score is independent of alignment length, an advantage over other structure similarity scores (Zhang and Skolnick, 2004). SID is the fraction of aligned residues that have the same amino acid identity. There are *N* * (*N* – 1)/2 unique pairs, equating to 278, 631 *S. cerevisiae* and 602, 253 *E. coli* protein pairs in ProteomeVis.

#### 2.1.3 Length and contact density

A protein’s length is the number of residues present in the PDB structure file. The contact density is the total number of residue-residue contacts in the PDB structure file, normalized by its length. Two residues are defined to be in contact if any non-hydrogen atom of the residue is within 4.5 Angstroms (Å) from another non-hydrogen atom of a non-adjacent residue (Bloom *et al.*, 2006; Zhou *et al.*, 2008). In Results 3.2, we also explore defining contacts as residue pairs having Cβ (C_β_ for glycine) distances <7.5 Å (Shakhnovich, 2006). The Biopython module (Cock *et al.*, 2009) is employed to parse PDB files and calculate length and contact density.

#### 2.1.4 Protein abundance

Protein abundance per cell was measured in Ghaemmaghami *et al.*, 2003 and Arike *et al.*, 2012 for *S. cerevisiae* and *E. coli* K-12 MG1655, respectively. Abundance data are downloaded from the Protein Abundances Across Organisms database (PaxDb) (Wang *et al.*, 2012). They cover 57% and 74% of the *S. cerevisiae* and *E. coli* K-12 proteomes. They are the largest and most accurate single-sourced datasets for their respective proteome (PaxDb accessed in November 2017). To allow for fair comparison between organisms, abundance data in ProteomeVis are presented as protein abundance per cell divided by the total protein abundance per cell for the respective organism.

#### 2.1.5 Protein-protein interaction

PPI partners are downloaded from the IntAct database (Orchard *et al.*, 2014) via the Bioservices Python module (Cokelaer *et al.*, 2013). IntAct has the largest number of experimentally determined PPIs among PPI databases (Szklarczyk and Jensen, 2015). Only interactions identified by affinity purification methods for *S. cerevisiae* S288c and *E. coli* K-12 are included. Affinity purification methods capture more stable interactions, rather than transient interactions that can vary greatly between independent experiments (Berggård *et al*., 2007).

#### 2.1.6 Dosage tolerance

Dosage tolerance was measured in Douglas *et al.*, 2012 for *S. cerevisiae* and in Kitagawa *et al.*, 2005 for *E. coli* K-12. Douglas *et al.*, 2012 reported dosage tolerance, called overexpression toxicity in their paper, as a logarithmic base 2 (log2) ratio between organismal growth when the gene of interest is overexpressed compared to that when the gene is expressed at wild-type levels. Dosage sensitive genes are defined as those with log2 ratios less than -1. Kitagawa *et al.*, 2005 reported dosage tolerance, called growth inhibitory effects in their paper, as an integer value of 1, 2, or 3. One signifies no growth, 2 signifies some growth and 3 signifies wild-type growth when the gene is overexpressed. All studied genes, regardless of their wild-type expressions, are overexpressed to the same elevated copy number. The most recent and least invasive dosage tolerance measurement for *S. cerevisiae* is from Douglas *et al.*, 2012. Kitagawa *et al.*, 2005 is the only known study that measured dosage tolerance for *E. coli*.

#### 2.1.7 Evolutionary rate

ER data for *S. cerevisiae* S288c and *E. coli* K-12 MG1655 are taken from Zhang and Yang, 2015. Briefly, orthologous proteins for *S. cerevisiae* S288c were identified in *Saccharomyces baynus* and for *E. coli* K-12 MG1655, in *Salmonella typhimurium* LT2. ER was calculated as the SID between orthologous proteins. This approximates nonsynonymous substitutions per nonsynonymous site (d_N_), a more popular metric for ER. Indeed, the correlation between ER from Zhang and Yang, 2015 and *d_N_* from Wall *et al.*, 2005 for *S. cerevisiae* is very strong - Spearman correlation=0.98, p-value<10^−300^ (Figure S1). We will add *d_N_* and *d_S_* data in future iterations of ProteomeVis.

### 2.2 Web app

The ProteomeVis web app can be accessed through any web browser at http://proteomevis.chem.harvard.edu/proteomevis/. To access the source code, refer to https://github.com/rrazban/proteomevis. The web framework of ProteomeVis is implemented using Python Django. The front-end is implemented in JavaScript with D3.js; the backend, SQLite. ProteomeVis is hosted on Amazon Web Services. Currently, users can only upload their data by altering the CSV files from which the database is created and deploying the web app locally.

The ProteomeVis web app layout comprises a top bar - called the Control Strip (Figure 1A), two left panels - Edge Filtering (1B) and Protein Chain Graph (PCG) (1C), one center panel - Protein Inspection (1D), and one right panel - Scatter Plot Matrix (SPloM) (1E).

#### 2.2.1 Control Strip

The Control Strip (Figure 1A) controls the overall data presented. From left to right, the user can (1) select which organism’s proteome to visualize (2) enter the TM-score and SID ranges that define a graph in the PCG panel (3) select the differentially colored property in the PCG panel in log10 units, except for dosage tolerance (no log transformation) (4) click the download button to obtain spreadsheets of the underlying data (5) click the help button for a quick tour of the web app. In Figure 1, the Control Strip is set up such that the *S. cerevisiae* proteome is visualized. Protein pairs with TM-score>0.5 and SID<0.25 are connected by an edge and nodes are differentially colored by degree in the PCG.

#### 2.2.2 Edge Filtering

The Edge Filtering panel (Figure 1B) displays the SID vs. TM-score scatter plot for all protein pairs in the designated proteome. As an alternative to entering values into the TM-score and SID fields in the Control Strip, users can define TM-score and SID ranges by brushing over the SID vs. TM-score plot. Each point corresponds to a potential edge in the PCG. All edges within the brushed region are rendered in the PCG panel below the Edge Filtering panel. In addition, datapoints colored green denote protein pairs that physically interact according to the IntAct database.

#### 2.2.3 Protein Chain Graph

Upon setting TM-score and SID ranges, nodes and edges appear in the PCG panel (Figure 1C). A node represents an individual protein, while

**Fig. 1:**
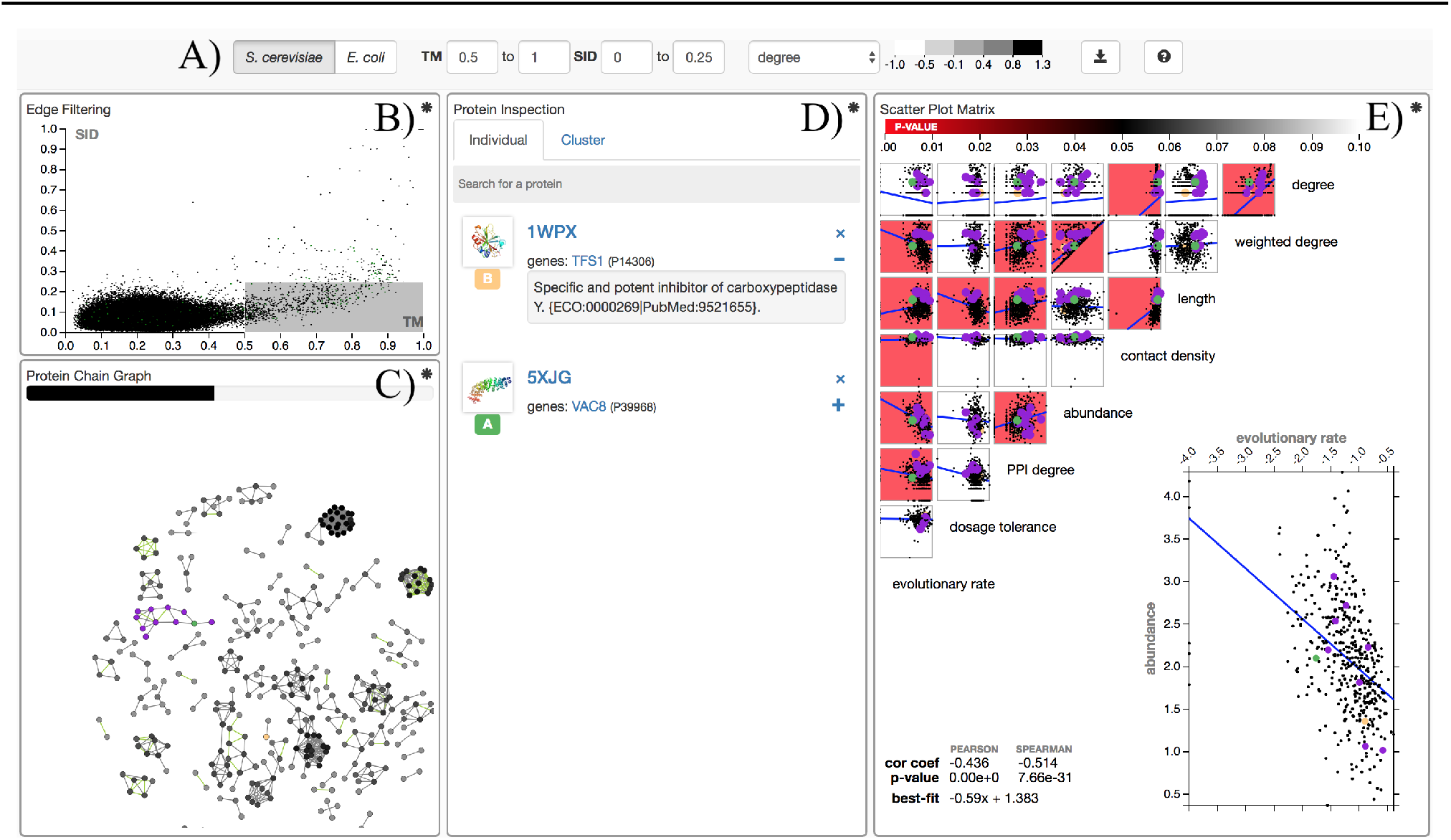
The ProteomeVis web app after exploration has occurred. Hovering over asterisks located in the top-right corner of each panel opens an informational window. Pressing the help button (question mark symbol) located to the right of the Control Strip (A) leads to a quick tour. Alphabetical labels (A-E) are added to each component of ProteomeVis in this figure to introduce the tool in the text.

an edge signifies that the two proteins it connects have a TM-score and SID within the user-defined range. Only nodes with at least one edge are shown in the PCG for visual clarity. The horizontal bar at the top of the panel allows users to gauge the fraction of nodes displayed (black). As in the Edge Filtering panel, edges colored green denote two physically interacting proteins. The node’s color is given by the gray scale shown in the Control Strip. In Figure 1, nodes are colored by degree. Nodes in small clusters tend to be light in color as they have low degrees, while nodes in large clusters are darker because they are typically connected to multiple other nodes. When the user hovers over a node, the protein structure and PDB identifier appear at the top left of the panel. If the user clicks on the node, detailed information about the protein appears in the Protein Inspection panel.

#### 2.2.4 Protein Inspection

The Protein Inspection panel (Figure 1D) invites users to examine the properties of any protein in ProteomeVis. Clicking on a node in the PCG or searching for a protein in the search bar populates the Individual tab of the Protein Inspection panel with the corresponding protein complex and its chains, PDB identifier, associated genes and UniProt accession numbers. Clicking the plus button provides information on the protein complex’ function as well as PubMed identifiers and Evidence codes (ECOs) (Chibucos *et al.*, 2014) for tracking references. The PDB identifier and gene name are clickable links to its PDB and UniProt web page, respectively. Clicking the image of the protein structure opens a window with the image enlarged and protein property values below it.

The Cluster tab in the Protein Inspection panel makes it easy for users to peruse all proteins in a connected component in the PCG (Figure S2). The cluster size is the number of nodes comprising the cluster in the PCG. The tab displays a histogram of cluster sizes, for all cluster sizes larger than 2. In the histogram, each cluster is represented by a square. The gray scale color of each square in the histogram denotes the average degree (default) or other property defined by the user in the Control Strip. All proteins belonging to a cluster can be displayed by clicking on the square representing the cluster, which displays a header, and then clicking this header.

Protein chains identified in the Individual and Cluster tabs are given a color, which allows them and data associated with them to be identified in the PCG and SPloM panels, respectively. In these tabs, the color is displayed in the protein chain letter underneath the image of the protein structure. In Figure 1,1WPX chain B is colored orange. Its representative node in the PCG panel and its associated datapoints in the SPloM panel are also orange. If the default color assigned is not appealing, the user can cycle through different colors by clicking the protein chain letter. For example, in Figure S2, all chains associated with the selected cluster are displayed in purple. In order to differentiate 5XJG chain A from other proteins in the cluster, the protein chain letter was clicked, yielding a green color (Figure 1D).

#### 2.2.5 Scatter Plot Matrix

The SPloM panel (Figure 1E) allows users to quickly identify statistically significant correlations between pairs of protein properties. ProteomeVis includes eight properties.

1. Degree: number of nodes sharing an edge with a node of interest in the PCG, e.g. 4 for the gray node in Figure S3A.
2. Weighted degree: the abundance of a protein of interest plus the sum of protein abundances that share an edge with the protein of interest in the PCG, e.g. 12 for the gray node in Figure S3.
3. Length: number of residues in the PDB structure.
4. Contact density: average number of contacts per residue in the PDB structure.
5. Abundance: normalized protein copy number per cell in parts per million.
6. PPI degree: number of stable protein-protein interaction partners.
7. Dosage tolerance: an organism’s level of insensitivity to protein overexpression.
8. Evolutionary rate: SID between orthologous proteins.

Not all proteins in ProteomeVis have recorded values for all of its properties. The last four in the list are obtained from external sources as described in Methods 2.1, and some have incomplete overlap with ProteomeVis (Figure S4).

Each of the 8*7/2 = 28 graphs in the panel, except for those involving dosage tolerance, is a log-log scatter plot of one trait against another. Values of 0 are approximated by an order of magnitude smaller than the second smallest possible value for the respective protein property. For example, the second smallest value for degree is 1, therefore 0 is approximated by 0.1 in the log scale. The scatter plots involving the last 6 properties in the list are static, i.e. no matter the TM-score and SID ranges chosen, the plots remain the same. Only plots involving degree and weighted degree change each time a new TM-score and SID selection is made, such that they always reflect the PCG. As described above, nodes with no edges, called orphans, are not displayed in the PCG. However, data associated with orphans are always shown in the SPloM, including their degree (i.e. zero, approximated by 0.1 in the log scale) and weighted degree (i.e. their abundance).

All plots in the SPloM have a background color determined by the p-value color bar at the top of the panel. Red corresponds to a p-value<0.01; black, p-value=0.05; and white, p-value>0.1. If the Spearman correlation coefficient is less than 0.15, no matter how small the p-value, the plot will be colored white. Results from correlation analysis can be obtained by hovering over a plot in the SPloM. Clicking on a specific scatter plot enlarges it in the bottom right corner of the SPloM panel for closer inspection and presents its corresponding correlation analysis results to its left. In Figure 1, the abundance vs. evolutionary rate plot is enlarged.

#### 2.2.6 Data download

At any point, users can download the data making up ProteomeVis by clicking the download button located on the far right of the Control Strip. As shown in Figure S5, there are three main options. First, users can download a spreadsheet of the individual proteins’ properties, selecting which of the eight properties they desire. Second, they can choose to download data on the relationships between protein pairs, either the edges that fulfill the set TM-score and SID ranges (“Edges in current range”) or all edges. Edge files include TM-scores and SIDs for each protein pair, as well as whether they form a PPI. Third, in the Correlations tab, users can download matrices of correlation analyses values presented in the SPloM panel.

In addition to downloading from the Control Strip, ProteomeVis offers a fourth option that enables users to download data associated with specific protein clusters. In the Cluster tab of the Protein Inspection panel, clicking on a cluster, hovering over its header line, and clicking the download icon that appears to the left, downloads the data. In Figure S2, the download button is not seen but if the user was to hover over the header line with their cursor, the download button would appear to the left of the 11 label.

We encourage users to download data to rigorously test any hypotheses. Simply scanning the SPloM panel for significant correlations is prone to the multiple testing problem and may lead to incorrect conclusions (Yoav and Hochberg, 1995).

**Fig. 2:**
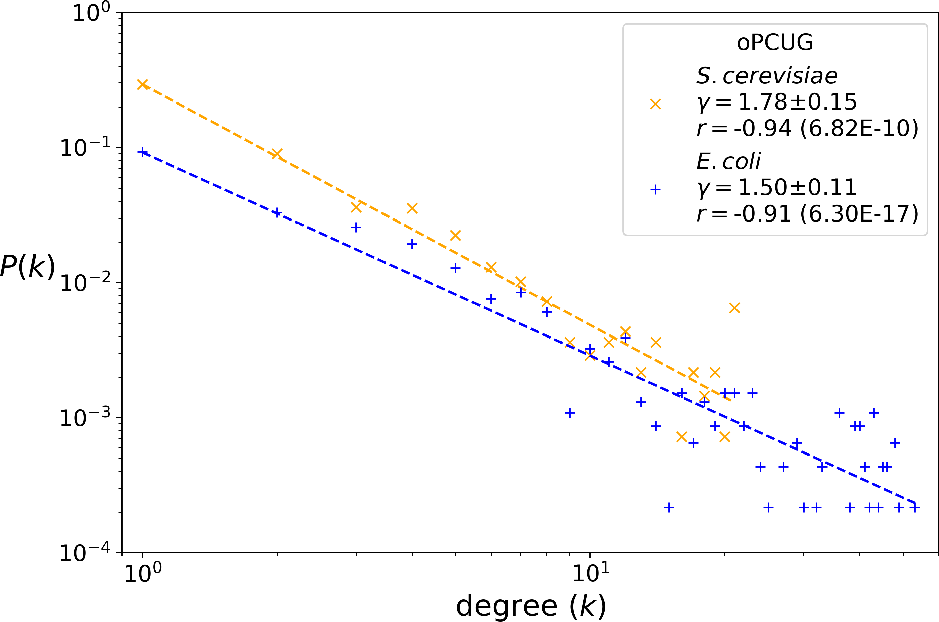
Degree distribution of the organismal protein chain universe graph (oPCUG) with its best-fit line on a log-log plot. The slope of the best-fit line (scaling constant, 7) with its standard error and the Pearson correlation coefficient (*r*) with its p-value in parenthesis are calculated by Python SciPy’s linear regression function. Only 21 *S. cerevisiae* datapoints are seen because its maximal degree is 20; for *E. coli*, its maximal degree is 52. All degrees are horizontally shifted by 1 unit to allow viewing of orphans (degree=0) and be consistent with Deeds *et al*., 2004 in evaluating the scaling constant.

## 3 Results

### 3.1 Degree distribution of the organismal protein chain universe graph

A characteristic behavior seen in the PDUG compiled across (Dokholyan *et al.*, 2002) as well as within organisms (oPDUGs) (Deeds *et al.*, 2004; Roland and Shakhnovich, 2007) is that they are scale-free, meaning that the probability to find a node with degree *k* scales as a power law of *k* with a scaling constant, γ: *P*(*k*) ~ *k*^−γ^. This behavior, especially for oPDUGs, provides support that protein structures evolutionarily diverge rather than converge. In Figure 2, ProteomeVis reproduces the scale-free behavior for the *S. cerevisiae* and *E. coli* oPCUGs. The oPCUGs are composed of protein pairs from the respective organism with TM-scores greater than 0.5 and SIDs less than 0.25. For an indepth discussion on threshold and cutoff values, and the scaling constant’s robustness, refer to Supplementary information 3.1a. To create Figure 2, the TM-score and SID ranges are entered in the Control Strip of the ProteomeVis web app and then the degree data is downloaded via option 1 as described in Methods 2.2.6.

The slope of the best-fit line in a log-log plot is the additive inverse of the scaling constant. Both organisms exhibit highly significant scale-free behavior, with Pearson correlation coefficients<–0.9 and corresponding p-values<10^−9^ (Figure 2). *E. coli’s* oPCUG scaling constant is 1.50 ± 0.11, whose range captures Deeds *et al.*, 2004’s measurement of 1.59 for *E. coli*’s oPDUG. We hypothesize that this agreement results from protein domains within protein chain pairs dominating alignments and leave it for a future study. *S. cerevisiae*’s scaling constant is 1.78 ± 0.15. No corresponding protein universe graph result is present in the literature for *S. cerevisiae.* Qian *et al.*, 2001 and Koonin *et al.*, 2002 demonstrated that the distribution of *S. cerevisiae* domains exhibiting the same structure classified by SCOP are scale-free. However, Dokholyan *et al.*, 2002 showed that this distribution is one of cluster sizes, making its scale-free behavior simply a result of random graphs and not indicative of any unique mechanism behind protein structure evolution.

Both organisms exhibit oPCUG scaling constants that fall within the 1.4- 1.9 range of scaling constants output by Big Bang model simulations (Dokholyan and Shakhnovich, 2001), described in the Introduction. This result has interesting implications for the generality of protein structure evolution. Although *S. cerevisiae* and *E. coli* belong to different kingdoms of life, their oPCUG scaling constants are captured by the Big Bang model, indicating that the mechanism behind protein structure evolution may be inherent to the last universal common ancestor.

### 3.2 Putative evolutionary rate determinants

Multiple studies have determined protein expression to be the strongest predictor of ER (Pál *et al.*, 2001; Drummond *et al.*, 2005, 2006; Drummond and Wilke, 2008; Zhang and Yang, 2015). Quantified by mRNA abundance, protein expression correlates negatively with ER. There has been continuous debate as to whether other properties impact ER. Figure 3 summarizes the correlations between ER and all five non-PCG properties in ProteomeVis, alongside their corresponding correlations in the literature, if available. The scatter plots from which correlation coefficients and p-values are calculated, can be seen in the SPloM panel of the ProteomeVis web app. Protein abundance has the largest magnitude correlation coefficient at -0.51 for *S. cerevisiae* and -0.46 for *E. coli*, with the lowest p-values (<10^−10^), in agreement with protein expression results. Previous studies have mostly focused on the correlation between mRNA abundance and ER because measuring protein abundance across the proteome was more laborious (Vogel and Marcotte, 2012). Just one other study examined the protein abundance-ER relationship in *S. cerevisiae*, and they reported a strong negative correlation (Drummond *et al.*, 2006). To our knowledge, no other study has performed this analysis for *E. coli*. Our results expand the scope of previous protein expression studies by demonstrating that a protein level property, protein abundance, strongly correlates with ER for both *S. cerevisiae* and *E. coli.* This result is nontrivial because the correlation between mRNA and protein abundance is not very strong (Greenbaum *et al.*, 2003; Vogel and Marcotte, 2012).

The second strongest ER determinant in ProteomeVis is PPI degree. In agreement with Fraser *et al.*, 2003, we report a significant negative correlation between PPI degree and ER (Figure 3). Fraser *et al.*, 2003 hypothesized that proteins with more interaction partners are under stronger negative selection to keep their sequences unchanged to avoid detrimental loss of interaction partners. Bloom and Adami, 2003 argued that the negative correlation between PPI degree and ER is confounded by protein expression. In the dataset that Fraser *et al.*, 2003 used, Bloom and Adami, 2003 found that proteins with more PPIs were more likely to have larger protein expression measurements. When controlling for protein expression by partial correlation analysis, Bloom and Adami, 2003 found no significant correlation between PPI degree and ER. To see if this correlation as revealed by ProteomeVis could be confounded by protein abundance, the PPI degree vs. abundance plot is explored in the SPloM panel. Indeed, there is a significant positive correlation between them in ProteomeVis - Spearman correlation=0.27, p-value<10^−10^, whose absolute value is approximately the same as that of the PPI degree vs. ER correlation - Spearman correlation=-0.27, p-value<10^−9^.

Focusing on the correlation between contact density and ER, ProteomeVis can explain previously conflicting reports for *S. cerevisiae.* Mathematical relationships predict that the number of sequences stably folding into a native structure is approximated by contact density (England and Shakhnovich, 2003; Choi *et al.*, 2017). Therefore, proteins with larger contact densities are expected to have larger ERs because more mutations can be accommodated while still stably folding. In agreement, Zhou *et al.*, 2008 reported a significant positive correlation between contact density and ER for *S. cerevisiae* and *E. coli.* However, Shakhnovich, 2006 found a corresponding correlation for *S. cerevisiae* that is significantly negative. In ProteomeVis, corresponding positive correlations are reported that is significant (p-value<10^−3^) for *S. cerevisiae* and slightly significant (p-value<0.01) for *E. coli.* To probe whether ProteomeVis’ consistency with Zhou *et al.*, 2008 is due to having the same contact definition, we define contacts as in Shakhnovich, 2006 (Methods 2.1.3). The corresponding correlation for *S. cerevisiae* becomes insignificant - Spearman correlation=0.02, p-value>0.05. Although we do not recapitulate the significant negative correlation found in Shakhnovich, 2006, we find that the significant positive correlation between contact density and ER in ProteomeVis is contingent on the employed contact definition.

As seen In Figure 3, all statistically significant correlations in ProteomeVis agree with those available in the literature, except for the correlation between length and ER for *E. coli*. Although ProteomeVis agrees with Zhou *et al.*, 2008 regarding correlations between contact density and ER, they contradict regarding the correlation between protein length and ER for *E. coli.* Zhou *et al.*, 2008 reported a significant positive correlation, while ProteomeVis reports an insignificant negative correlation for *E. coli*. Methodological differences may still explain the discrepancy. Length could be more sensitive to methodology than contact density because contact density is normalized while length is an absolute number. Inaccuracies in the number of contacts and length due to the identified PDB structure being poor, could partially cancel out when dividing the two numbers to obtain contact density. Zhou *et al.*, 2008 employed a SID threshold of 0.4, compared to 0.9 in ProteomeVis, and did not filter based on length to identify structures in the respective organism’s proteome. In spite of methodological differences, the length vs. ER correlation for *S. cerevisiae* in ProteomeVis and Zhou *et al.*, 2008 are consistent. Further studies are required to elucidate why differences in protein structure identification lead to an inconsistent length-ER correlation in *E. coli* but not *S. cerevisiae.*

## 4 Conclusion

Proteomic data visualization tools are increasingly developed to facilitate understanding of complex and interwoven experimental data (Vizcaíno *et al.*, 2015). Here, we present a novel web app called ProteomeVis, accessible at http://proteomevis.chem.harvard.edu/proteomevis/, to study protein structure and sequence evolution simultaneously across organisms’ proteomes. Data downloaded from three published papers, programmatically accessed from four databases and generated by running three software packages are organized into ProteomeVis’ four panels. Interacting with the panels gives users quick insight into the underlying data. Once more detailed hypotheses are formulated throughout their web sessions, users can freely download the data to further probe on their own.

In this paper, we applied ProteomeVis to investigate two important problems in evolutionary protein biology: the scale-free degree distribution of oPCUGs and the correlations between ER and protein properties. These two topics span time and length scales in protein evolution, and we successfully recapitulated and extended previous studies. We report the previously unknown scaling constant for *S. cerevisiae*’s protein universe graph. Interestingly, its oPCUG scaling constant fits the range predicted by the Big Bang model, which had only been applied to prokaryotic organisms (Dokholyan *et al.*, 2002; Deeds *et al.*, 2004). This observation points to the mechanism of protein structure evolution being common among organisms since the last universal common ancestor. In our studies of potential ER determinants, protein abundance is the strongest correlate among properties in ProteomeVis. Previously, the importance of mRNA abundance had been highlighted (Zhang and Yang, 2015), but our work joins only one other study to demonstrate the importance of protein abundance influencing ER (Drummond *et al*., 2006).

**Fig. 3:**
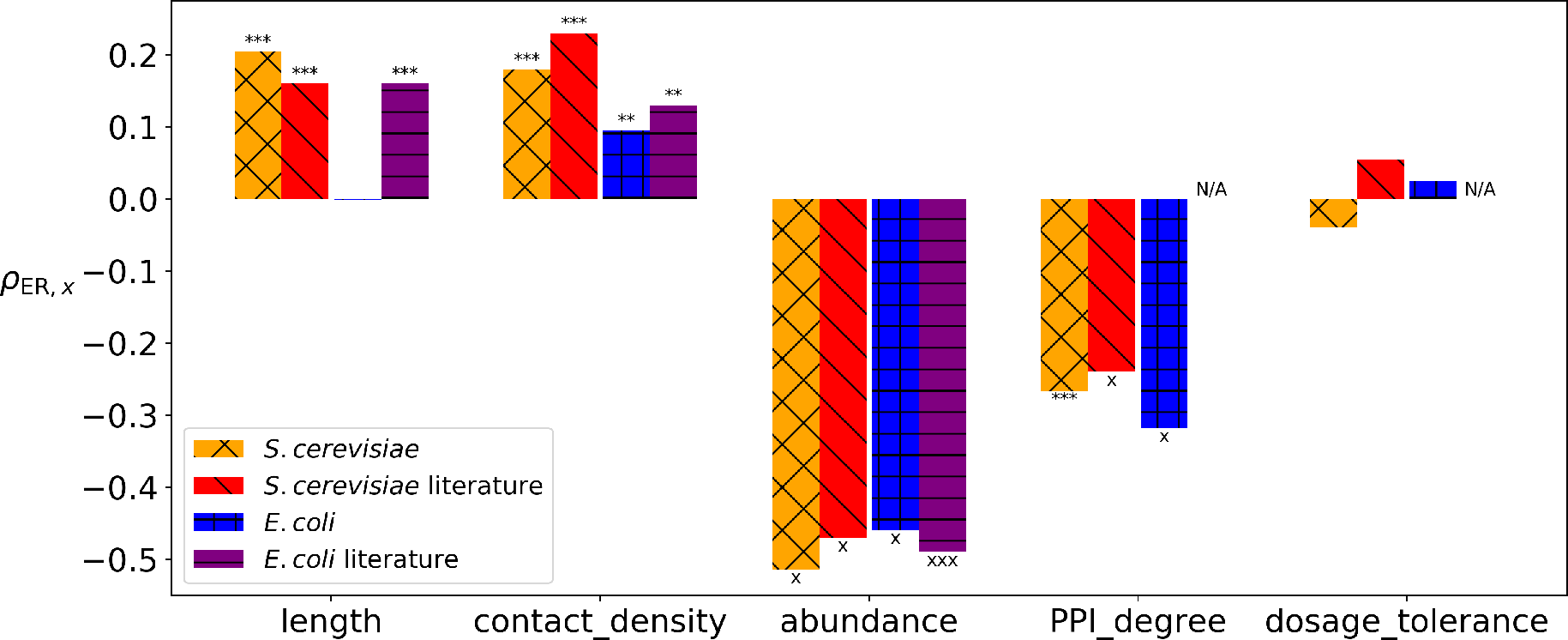
Spearman correlations (ρ) between evolutionary rate (ER) and five protein properties (x) in ProteomeVis and the literature. X marks and asterisks above or below bar plots denote p-value ranges: xxxΞ p-value < 10^−100^, xx< 10^−50^, x< 10^−10^, ***< 0.001, **< 0.01, *< 0.05. Length (Zhou *et al*., 2008), contact density (Zhou *et al.*, 2008) and abundance (mRNA) (Zhang and Yang, 2015) correlations with evolutionary rate in the literature for both organisms are presented. PPI degree and dosage tolerance correlations with evolutionary rate in the literature for *S. cerevisiae* are from Fraser *et al*., 2003 and Vavouri *et al.*, 2009, respectively; for *E. coli*, not available (N/A) as of November 2017.

ProteomeVis captures 12% of the *S. cerevisiae* and 25% of the *E. coli* proteomes as of November 2017. These percentages will grow as monthly updates are performed to incorporate newly deposited PDB structures. We plan to further expand ProteomeVis in future iterations by including more proteomes, such as those from *Homo sapiens* and *Mus musculus*, and adding genomic data, such as *d_N_* and *d_s_*.

## Acknowledgements

RMR acknowledges fruitful email communications with Professor Brenda Andrews about dosage tolerance data in Sopko *et al.*, 2006, and Professor Jianzhi Zhang and Professor Jian-Rong Yang about evolutionary rate data in Zhang and Yang, 2015. RMR would like to thank Will Jacobs for helpful discussions surrounding data curation. All figures in the text and Supplementary information, except for those of the ProteomeVis web app and Figure S3, were created using Python’s Matplotlib package.

## Funding

This work has been supported by National Institutes of Health [2R01GM068670-13 to E.I.S.]; and the National Science Foundation Graduate Research Fellowships Program [2013139945 to A.I.G.].

